# Morphogenomic description of *Cranifera cranifera* (Chitwood, 1932) Kloss, 1960 from captive *Blaptica dubia* Serville, 1838 cockroach

**DOI:** 10.64898/2026.07.17.739199

**Authors:** Jans Morffe, Nadège Guiglielmoni, Nadim Wassey, Karim Gueddach, Anja Schuster, Kerstin Becker, Jens Bast, Philipp H. Schiffer, Oleksandr Holovachov

## Abstract

Nematodes of the superfamily Thelastomatoidea are found in the digestive system of various arthropods, feeding on their host microbiome. They are sometimes considered to be ecologically intermediate forms between free-living rhabditids and parasitic Spirurina, while phylogenetically they are nested within the latter. In addition to new morphological data on the male morphology, this manuscript presents the first nuclear genome assembly of a thelastomatid species, *Cranifera cranifera,* using long-read sequencing approach, making a total of three nuclear genomes available for superfamilyThelastomatoidea. The *C. cranifera* nuclear genome assembly presented here is 246 Mb long, consists of 7563 contigs, has an N50 of 43 kb and includes 94% of the BUSCO *nematoda_odb12* genes. The mitochondrial genome is 24646 bases long, includes a complete set of protein coding, rRNA and tRNA genes, and a repetitive region 9731 bases long, which includes multiple copies of tRNA-Asn(gtt) and tRNA-Lys(ttt). The nuclear assembly also contained two sequence variants of the 28S rRNA and two of the 18S rRNA genes, highlighting the presence of intragenomic variation within rRNA operon. The newly generated assemblies (nuclear and organelle) will add to a growing body of genomic resources for underrepresented and understudied animal parasitic nematodes from the Clade 3, enabling comprehensive studies in their phylogeny and trait evolution in the future.

## Introduction

The genus *Cranifera* Kloss, 1960 belongs to a group of arthropod pinworms (superfamily Thelastomatoidea), a heterogeneous assemblage of Clade 3 nematodes found in the digestive system of diplopods, cockroaches, mole crickets, adult hydrophilid and passalid beetles, larvae of scarab beetles and crane flies (Carreno, 2014). Its type species was originally described under the name *Leidynema cranifera* Chitwood, 1932 from the hindgut of the cockroaches *Blaberus craniifer* Burmeister, 1838 and *B. atropos* (Stoll, 1813) from Florida, USA (Chitwood, 1932), and was later transferred into a genus of its own based on the absence of a diverticulum in the anterior portion of the intestine (Kloss, 1960). Females of *Cranifera* are characterized by the presence of a short first cephalic annule, unarmed cervical cuticle, corpus with a cylindrical anterior half and a sub-cylindrical, wider posterior half and didelphic genital tract (Adamson, 1992; Morffe et al., 2022). Males possess a spine-like caudal appendage, arising dorsally; three pairs of copulatory papillae arranged as one pre-cloacal and two post-cloacal (one adanal pair and another near the base of the tail appendage) and a single spicule (Chitwood, 1932).

Besides the type species of the genus, three additional species have been described: (a) *C. chitwoodi* Kloss, 1966 from the cockroach *Hormetica scrobiculata* Burmeister, 1838 from Brazil, (b) *C. mexicana* Coy & García, 1995 from death’s head cockroach *B. craniifer* from Mexico, and (c) *C. robustum* Camino & Achinelly, 2012 from the larvae of an Argentinean scarab *Cyclocephala signaticollis* Burmeister, 1847 from Argentina (Kloss, 1966; Coy and García, 1995; Camino and Achinelly, 2012).

Since Chitwood (1932) description *C. cranifera* has been recorded in several wild and laboratory reared blaberid cockroaches from Europe and The Americas, namely captive colonies of death’s head cockroach *B. craniifer* and the dubia roach *Blaptica dubia* Serville, 1838 from Germany (Leibersperger, 1960), laboratory colonies of *B. dubia* from Russia (Spiridonov and Guzeeva, 2009), as well as wild populations of the peppered cockroach *Archimandrita tessellata* Rehn, 1903 from Costa Rica, and the discoid cockroach *Blaberus discoidalis* Serville, 1838 from Cuba (Morffe et al., 2022). Males of *C. cranifera* were found and described only by Chitwood (1932) and Leibersperger (1960), however some taxonomically valuable features of male morphology were not sufficiently described. In this publication a population of *C. cranifera* from laboratory reared colonies of the dubia roach was used to provide a detailed redescription of the males of the species as well as to generate the nuclear and mitochondrial genomes using long-read sequencing data. We use this opportunity to propose a new term “morphogenomic species description”, in order to highlight/distinguish papers combining full morphology-based species descriptions together with detailed description of the genome of a species (already published examples include Slos et al., 2017; Stevens et al., 2019; Pettrich et al., 2026) and contrast them from: (1) traditional morphology-based (sometimes but not always accompanied by molecular data represented by few loci) species descriptions; (2) genome notes focusing solely on descriptions of genomes of known species and providing the briefest morphological data in the shape of voucher pictures, if any at all (examples include Röseler et al., 2025; https://www.darwintreeoflife.org/genomes/genome-notes/); and (3) genome-based species descriptions that include minimal or partial morphological information, often relegated to supplementary data (Dulovic et al., 2020; Launay et al., 2020; Devi et al., 2025).

## Material and methods

### Processing of the hosts and nematodes

Specimens of *Blaptica dubia* Serville, 1838 (Blattodea: Blaberidae) from a captive colony were killed by decapitation and immediately dissected using longitudinal incisions in both abdominal pleural membranes. Intestines were withdrawn from the body and dissected in dishes with 0.9% NaCl physiological solution. Nematodes for morphological studies were killed with hot 0.9% NaCl (70°C) and fixed in 4% phosphate buffered formalin. For light microscopy the nematodes were transferred to anhydrous glycerin via the slow evaporation method (Seinhorst, 1959) and mounted on permanent slides in glycerin with paraffin wax as support for the coverslip. The edges of the coverslips were sealed with nail polish. Several specimens were pooled for genomic studies and directly fixed in 1X DNA/RNA Shield™ (Zymo Research, California, USA) and stored at −20°C until DNA extraction.

### Morphological and morphometric studies

Nematodes were measured with the aid of a drawing tube attached to a Nikon Eclipse Ni microscope (Nikon, Tokyo, Japan). All curved structures were measured along the curved median line. De Man’s indices a, b, c and V% were calculated. Variables are shown as the range followed by the mean plus standard deviation in parentheses, the number of measurements is also given when it is different from the total number of measured specimens. Micrographs were generated with a Sony ɑ6400 digital camera attached to a Nikon Eclipse 80i compound microscope (Nikon, Tokyo, Japan). Line drawings were made with the aid of a drawing tube. Scale bars of all figures are given in micrometers. In total, 19 females (SMNH 231508–231516) and 5 males (SMNH-231517–231518) were examined for morphology.

### SEM studies

Nematodes were post-fixed overnight with 2% glutaraldehyde in 0.1 M phosphate buffer (pH 7.0). They were dehydrated through a graded ethanol series (30%, 50%, 70%, 90%, 95%, 100% × 2, 30 min in each); transferred to hexamethyldisilazane (HMDS) through a graded series of 25%, 50% and 75% of HMDS in 100% ethanol (30 min in each); washed in pure HMDS for 30 min; and finally were kept overnight in a fume hood until total evaporation of the HMDS. Nematodes were then mounted on double sided conductive adhesive carbon tabs on a stub, sputter coated with gold using an AGB7234 high resolution sputter coater (Agar Scientific, Rotherham, UK), and observed with a Quanta FEG 650 scanning electron microscope (Thermo Fisher Scientific, MA, USA) at 10 kV accelerating voltage.

### Genome sequencing

Nematodes preserved in Zymo DNA/RNA Shield solution were disrupted manually with a pestle while frozen and high molecular weight DNA was extracted using the Zymo *Quick*-DNA HMW MagBead kit (Zymo Research, California, USA) following manufacturer’s protocol. Due to low concentration of extracted DNA, ultra-low-input (ULI) library was prepared with the *SMRTbell prep kit 3.0 Kit* (https://www.pacb.com/wp-content/uploads/Procedure-checklist-Amplifying-genomic-DNA-for-SMRTbell-library-preparation-and-HiFi-sequencing.pdf, Pacific Biosciences, CA, USA). Whole genome amplification was performed with the Universal Adapter System (Twist Bioscience, CA, USA) and the KOD Xtreme™ Hot Start DNA polymerase (Merck, Germany). Library underwent quality control (Fragment Analyzer, Agilent, CA, USA) and size selection (Pippin HT, Sage Science, MA, USA) in order to quantify and eliminate shorter DNA fragments, with the final target fragment size of 10-20 kilobases. Library was pooled and sequenced on a Revio system (Pacific Biosciences, CA, USA). PacBio generated BAM files were converted into FASTQ format using *samtools* (Danecek et al., 2021). usegalaxy.eu (The Galaxy Community, 2026) installation of *fastplong* (Chen 2023) was used to remove sequencing adapters, trim 5’ and 3’ sequence ends and remove sequences with average *phred* quality less than 20. usegalaxy.eu (The Galaxy Community, 2026) installation of *NanoPlot* (De Coster and Rademakers, 2023) was used to assess the quality of the filtered raw read dataset.

### Nuclear genome assembly

Draft genomes were assembled with *Flye* (Kolmogorov et al., 2019), *Hifiasm* (Cheng et al., 2024) and *PECAT* (Nie et al., 2024) using *<u>usegalaxy.eu</u>* (The Galaxy Community, 2026). Assembly mode (--pacbio-corr, –pacbio-raw, –pacbio-hifi), assembly type (regular or metagenomic) and coverage for initial assembly were varied in case of assembly with *Flye*. For *Hifiasm*, genome size estimation was left blank or pre-set to 100 Mb, while for *PECAT* both PacBio sequencing platforms (hifi and clr) were tested. Quantitative completeness of draft assemblies was assessed using *BUSCO v.6.0.0* comparison against the *nematoda_odb12* reference dataset consisting of 596 universal single-copy orthologous genes (Tegenfeldt et al., 2025). *barrnap* (Galaxy version 1.2.2) on the usegalaxy.eu server (equivalent to standalone version 0.9, https://github.com/tseemann/barrnap) was used to mine both assemblies for Eukaryotic rRNA in order to confirm the identification of a target species, and to check for presence of contaminants, in particular for the host DNA. Two draft assemblies were selected for genome reconciliation: standard assembly generated with *Flye* using “–pacbio-raw” mode under offered highest N50, while metagenomic assembly generated with *Flye* using –pacbio-hifi mode had the highest *BUSCO* completeness, although the differences were minor.

Completeness and contamination level of both draft assemblies were further evaluated using *BlobToolKit* (Challis et al., 2020; https://www.sanger.ac.uk/tool/blobtoolkit/), and contigs identified as “chordata” and “arthropoda”, as well as all non-metazoan contigs were removed. *quickmerge* (Chakraborty et al., 2016) was used to reconcile both assemblies using either one as a query or a reference. Haplotigs (variants of the same gene originating from different alleles in the population) were purged from both reconciled assemblies using *redundans* (Pryszcz and Gabaldón, 2016), since *purge_dups* (Guan et al., 2020) was not able to handle population-level heterozygosity. Purged assemblies were again evaluated with *BUSCO v.6.0.0* by comparing them against the four most current reference datasets: *nematoda_odb12* (596 single copy-orthologs)*, nematoda_odb12.1* (1477 single-copy orthologs)*, chromadorea_odb12* (1126 single-copy orthologs) and *chromadorea_odb12.1* (2034 single-copy orthologs) (Tegenfeldt et al., 2025) and the final chosen assembly was visualised using *BlobToolKit* (Challis et al., 2020). Annotation of protein coding genes was not performed due to absence of RNAseq data.

### Mitochondrial genome assembly

A first attempt to assemble a mitochondrial genome was done with *MitoHifi* pipeline (Laslett and Canbäck, 2008; Allio et al., 2020; Uliano-Silva et al., 2023) using the mitochondrial genome of *Hammerschmidtiella* sp. (KY399989.1) as a reference and two alternative *Hifiasm*-based assemblies as an input. This produced two consensus contigs, 24645 and 35654 bases long. The discrepancies in length were caused by mis-assembly and not by collapsed/uncollapsed repetitive regions, requiring the use of the following alternative approach. Nucleotide sequences of nematode mitochondrial genes were downloaded from the https://github.com/WormsEtAl/Nematode-Mitochondrial-Database (Gendron et al., 2023) and converted into a *BLAST*-compatible nucleotide mitochondrial reference database with the usegalaxy.eu (The Galaxy Community, 2026) installation of a *NCBI BLAST+ makeblastdb* (Altschul et al. 1990). Amino acid sequences of protein-coding genes were extracted from the mitochondrial genomes of 52 nematode species across the entire phylum downloaded from the NCBI GenBank, and converted into a *Diamond*-compatible amino-acid mitochondrial reference database with the usegalaxy.eu (The Galaxy Community, 2026) installation of *Diamond makedb* (Buchfink et al., 2021). *blastn* (Altschul et al., 1990) with E-value cutoff 0.001 was used to search the filtered raw reads against the nucleotide mitochondrial reference database, while *Diamond blastx* (Buchfink et al., 2021) set to invertebrate mitochondrial code and E-value cutoff 0.001 was used to search the filtered raw reads against the amino acid mitochondrial reference database. *BBedit* text editor was used to remove duplicate hits, and create a list of read sequence headers for the *seqkit* (Shen et al., 2024), which was used to pull relevant reads from the filtered raw reads file. Three input datasets, one based on *blastn* hits, one based on *diamond blastx* hits and one using combined hits were used as an input for the mitogenome assembly with usegalaxy.eu (The Galaxy Community, 2026) installation of *Canu* (Koren et al., 2017). Multiple assemblies were generated using the following constant settings: technology = PacBio; maximum corrected overlap mismatch = 0.2; target coverage for corrected reads = 10; low coverage depth = 5; estimated genome size = 25k. Four combinations of minimum read length / minimum overlap were used in parallel: 1000/500, 2000/1000, 3000/1500 and 4000/2000. *Mitos2* (Bernt et al., 2013) was used to extract COX1 from several randomly selected assemblies with the following settings: genetic code = Invertebrate (5), reference data = refseq63m. This COX1 sequence was then identified using NCBI blastn against GenBank core nucleotide database (excluding uncultured and environmental sample sequences). Same COX1 sequence was used to positively identify remaining assembled contigs as belonging to the same target species, orient them in the same direction, align and circularise using *AliView* (Larsson, 2014). Draft consensus assembly was created by combining all aligned contigs (circularised and partial) using majority rule approach. *minimap2* (Li, 2021) was used to map raw reads against draft assemblies using *map-hifi* setting while *pilon* (Walker et al., 2014) was used to improve draft consensus assembly based on *minimap2* output. *Tandem repeats finder* (Benson, 1999) and Claude AI by Anthropic were both used to identify repetitive regions and assembly artefacts in the final assembly. usegalaxy.eu (The Galaxy Community, 2026) installations of *Mitos2* (Bernt et al., 2013) and *Aragorn* (Laslett et al., 2004) were used to annotate final mitochondrial genome assembly, with additional manual checks aided by Claude AI by Anthropic using following genomes as references: *Aspiculuris tetraptera* (KT764937), *Enterobius vermicularis* (NC056632), *Hammerschmidtiella* sp. (KY399989), *Oxyuris equi* (NC027190), *Passalurus ambiguus* (NC028345), *Ruizia karukerae* (MF509850 and MF509851), *Rhigonema thysanophora* (NC024020), *Spauligodon atlanticus* (PX608399), *Syphacia muris* (AP017697), *S. obvelata* (KT900946), *Travassosinema* sp. (PX431499 and PX431500), *Wellcomia compar* (MW059037) and *W. siamensis* (NC016129). *samtools* (Danecek et al., 2021) was used to estimate the read coverage from the *minimap2* output while custom script (written by Claude AI by Anthropic on 24 May 2026) was used to normalize read depth values to get scores between 0-1. *Proksee* (Grant et al., 2023) was used to visualise the final assembly.

### Intragenomic variation of rRNA

*barrnap* (Galaxy version 1.2.2) on the usegalaxy.eu server (equivalent to standalone version 0.9, https://github.com/tseemann/barrnap) was used to mine ribosomal RNA genes from the final assembly. Several 28S rRNA and 18S rRNA sequences (partial fragments) were retrieved from GenBank and used in comparative/phylogenetic analysis. *Severianoia blapticola* Guzeeva, 2009 and *Travasossinema viatorum* Morffe, García & Hasegawa, 2023 were included as outgroups for the 28S rRNA and 18S rRNA analyses, respectively. The former species was selected for the 28S rRNA analysis since it is the closest sister group of *Cranifera*, despite having been considered a *species inquirenda* by Malysheva et al. (2020). Multiple sequence alignments were made using MUSCLE (Edgar, 2004) with the default parameters as implemented in MEGA12 (Kumar et al., 2024). MEGA12 was also used to identify the optimal model of evolution for the data set (GTR+I for the 28S rRNA and GTR+G+I for the 18S rRNA datasets) following the corrected Akaike Information Criterion (AICc) and to construct phylogenetic trees based on the Maximum Likelihood (ML) method. Nodal support was inferred by bootstrap analysis using 1,000 iterations. Pairwise distances were calculated for the 28S rRNA and 18S rRNA sequences of *C. cranifera* with MEGA12 using the Kimura 2-parameter model (Kimura, 1980).

### Use of Artificial Intelligence Tools

Claude AI by Anthropic was used to search and summarise published reference data, it was not used for writing the manuscript or for data analysis except for cases specifically described above.

## Results

### Additional data on male morphology

Body smaller and more slender than that of females (Table 1), slightly robust and fusiform, widening from the anterior end with maximum width at ca. first third of the body, then narrowing towards cloaca (Figure 1C). Anterior end of body truncated, posterior end ventrally curved, blunt with a dorsally located tail appendage. Lateral alae well-developed from the level of the last third of the isthmus to a very short distance (ca. 2 μm) before the level of the first pre-cloacal pair of papilliform sensilla (Figure 2D). Cephalic cap ca. 10 μm in diameter, 3 μm length, truncated, with smooth cuticle (Figure 2E). Cervical cuticle unarmed, presenting annuli from base of cephalic cap that extend towards posterior end of body, ending ventrally and laterally at level of the pre-cloacal pair of papilliform sensilla, then continuing dorsally to the level of the second post-cloacal pair of papilliform sensilla. The annuli from the cervical region are fine and at level of the isthmus become gradually wider towards the posterior end of body. Annuli of the last two thirds of the body ornamented with tubercles that form a single row across each annule (Figure 2C). Tubercles more prominent towards the posterior part of body. Mouth surrounded by a fine, wrinkled cuticular cordon followed by a loose, veliform labial ring anchored bilaterally to the cuticle via a frenulum-like structure (Figure 2F). Amphids lateral, slit-like, located in prominences of the labial ring (Figure 2F). Buccal capsule very short. Oesophagus consisting of a comparatively short, muscular, clavate corpus, well set-off from the long isthmus (Figure 1A). Anterior part of isthmus wider at the joint with the base of corpus then its diameter decreases towards the spherical basal bulb. Valve-plate of basal bulb well-developed. Intestine simple, sub-rectilinear. Nerve ring located at level of posterior third of isthmus. Excretory pore ventral, located at ca. 1.5–2 corresponding body-widths posterior the basal bulb (Figure 1A). Monorchic. Testis ventral, reflexed just posterior to the excretory pore, flexure ca. one corresponding body-width long. Drop-like spermatids, ca. 8 μm length. Cloaca located in a prominent cloacal cone. Spicule single, sclerotized, almost straight, shaft spindle-shaped, capitulum rounded, ventrally bent, and distal tip sharp (Figure 1D). Eight copulatory papilliform sensilla present, two pre-cloacal and six post-cloacal (Figure 1B, Figure 2D). Pre-cloacal pair sub-ventral, consisting of large, prominent, mammiform papillae, located at ca. 10 μm anterior to the level of cloaca. First pair of post-cloacal papillae smaller, sub-ventral, ad-cloacal. Second pair of post-cloacal papillae large and mammiform, sub-dorsal, near the base of the tail appendage. Third pair of post-cloacal papillae ventral, consisting of minute papillae located a level of proximal fifth of the tail appendage.

**Figure 1.**
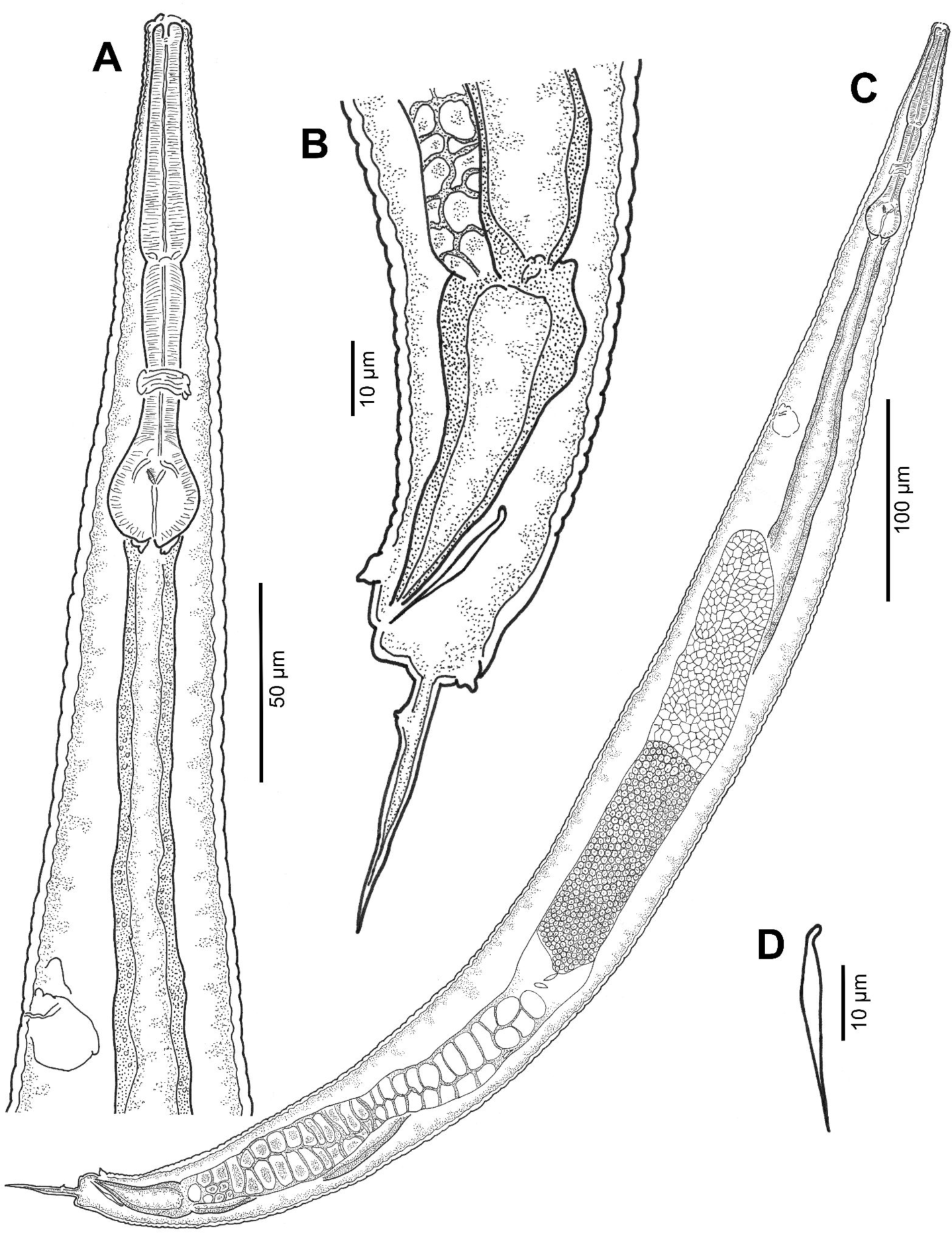
Line drawings of a male of *Cranifera cranifera* (Chitwood, 1932). A: Anterior body end showing pharyngeal region and excretory pore (ventral side to the left); B: Posterior body end showing cloaca, spicule, genital sensilla and tail (ventral side to the left); C: Entire male (ventral side to the left); D: Spicule (ventral to the right).

**Figure 2.**
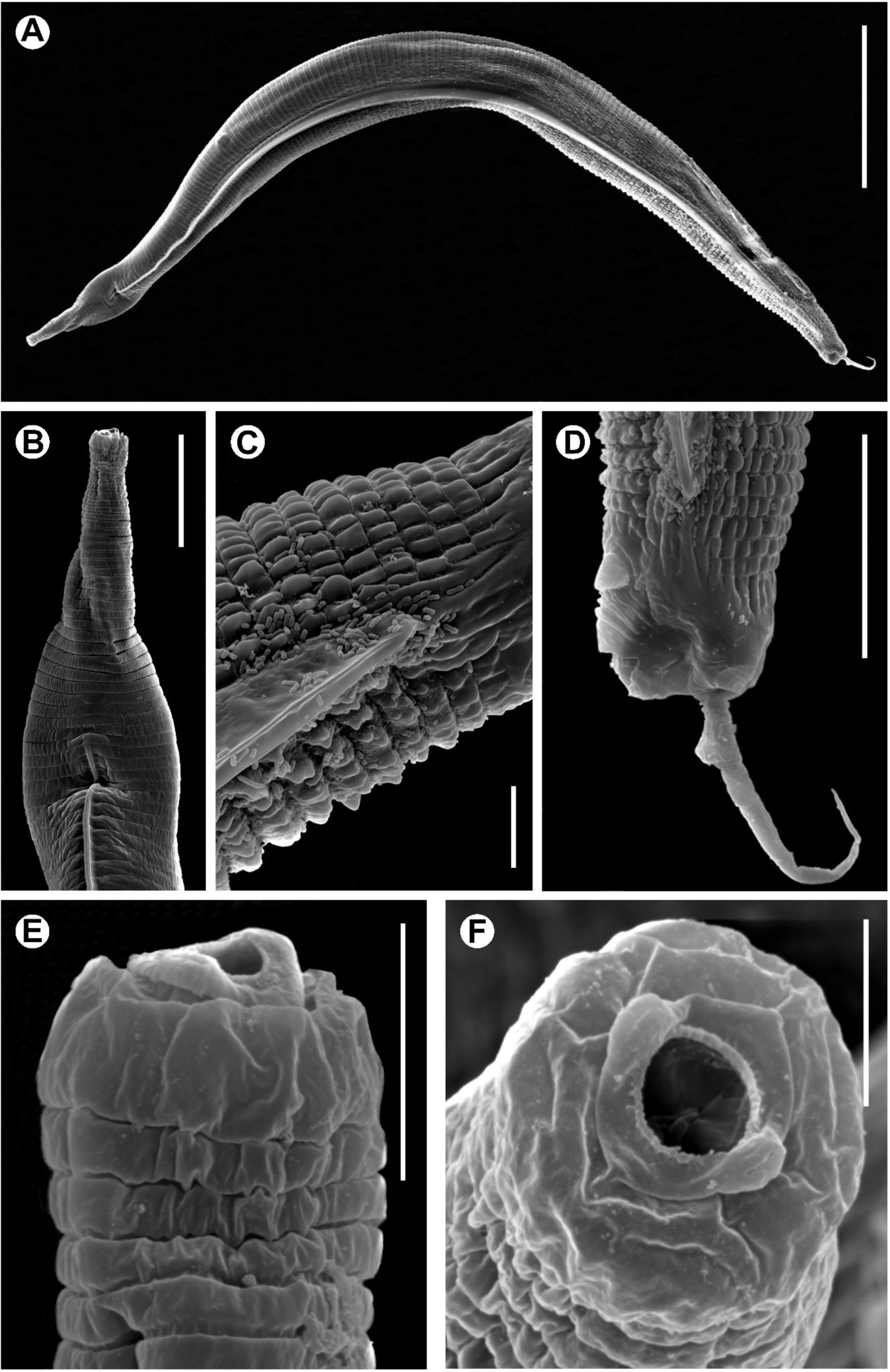
SEM micrographs of a male of *Cranifera cranifera* (Chitwood, 1932). A: Entire male (ventral side to the bottom); B: Anterior body end showing beginning of lateral alae (ventral side to the left); C: Posterior region showing the end of lateral alae and the ornamentations of the cuticle (ventral side to the bottom); D: Posterior body end showing cloaca, end of lateral alae, genital sensilla and tail (ventral side to the left); E: Labial region; F: Labial region, *en face* view. Scale bars: A = 100 *μ*m. B, D = 20 *μ*m. C, E = 5 *μ*m. F = 3 *μ*m.

**Table 1.**
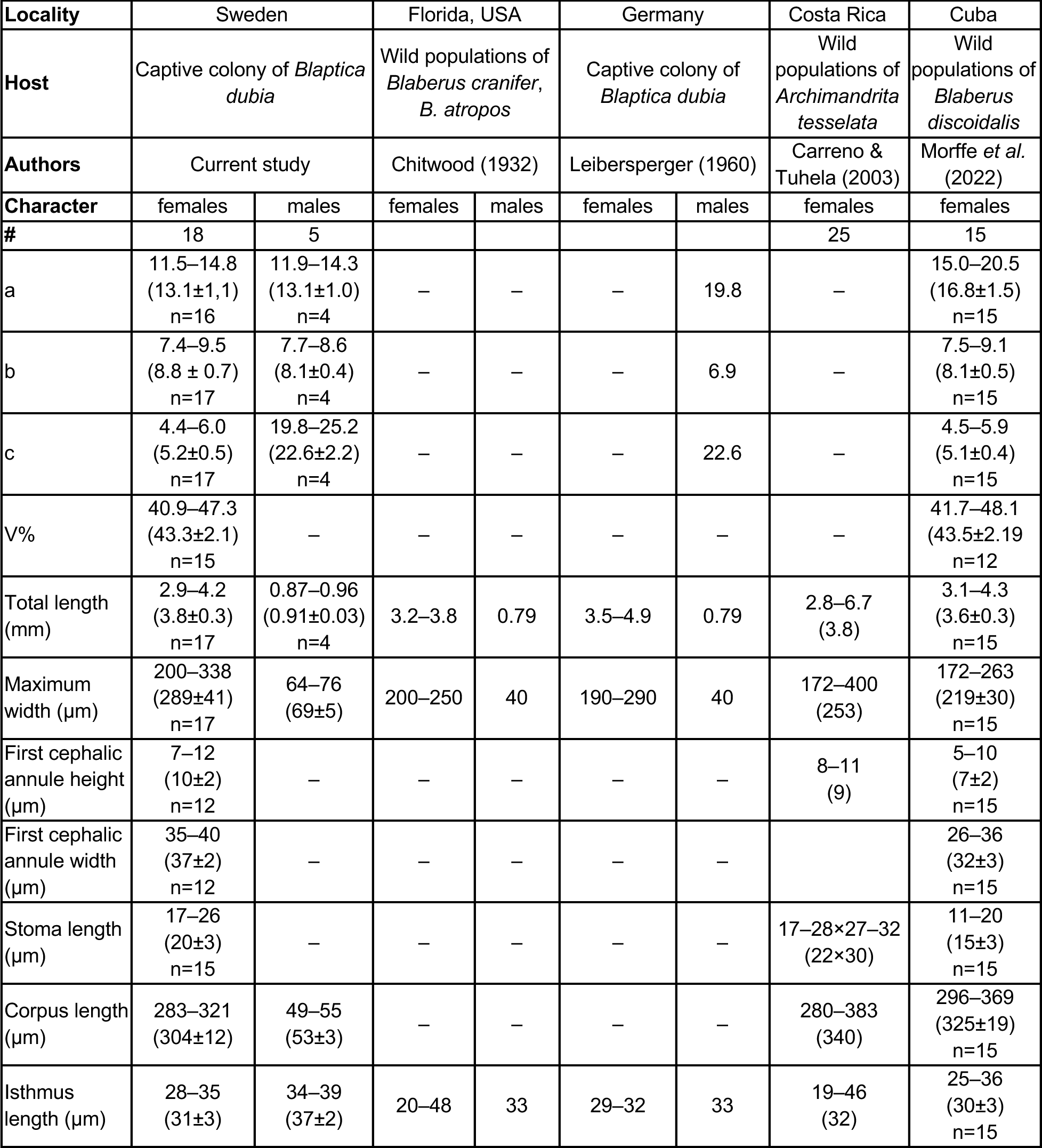
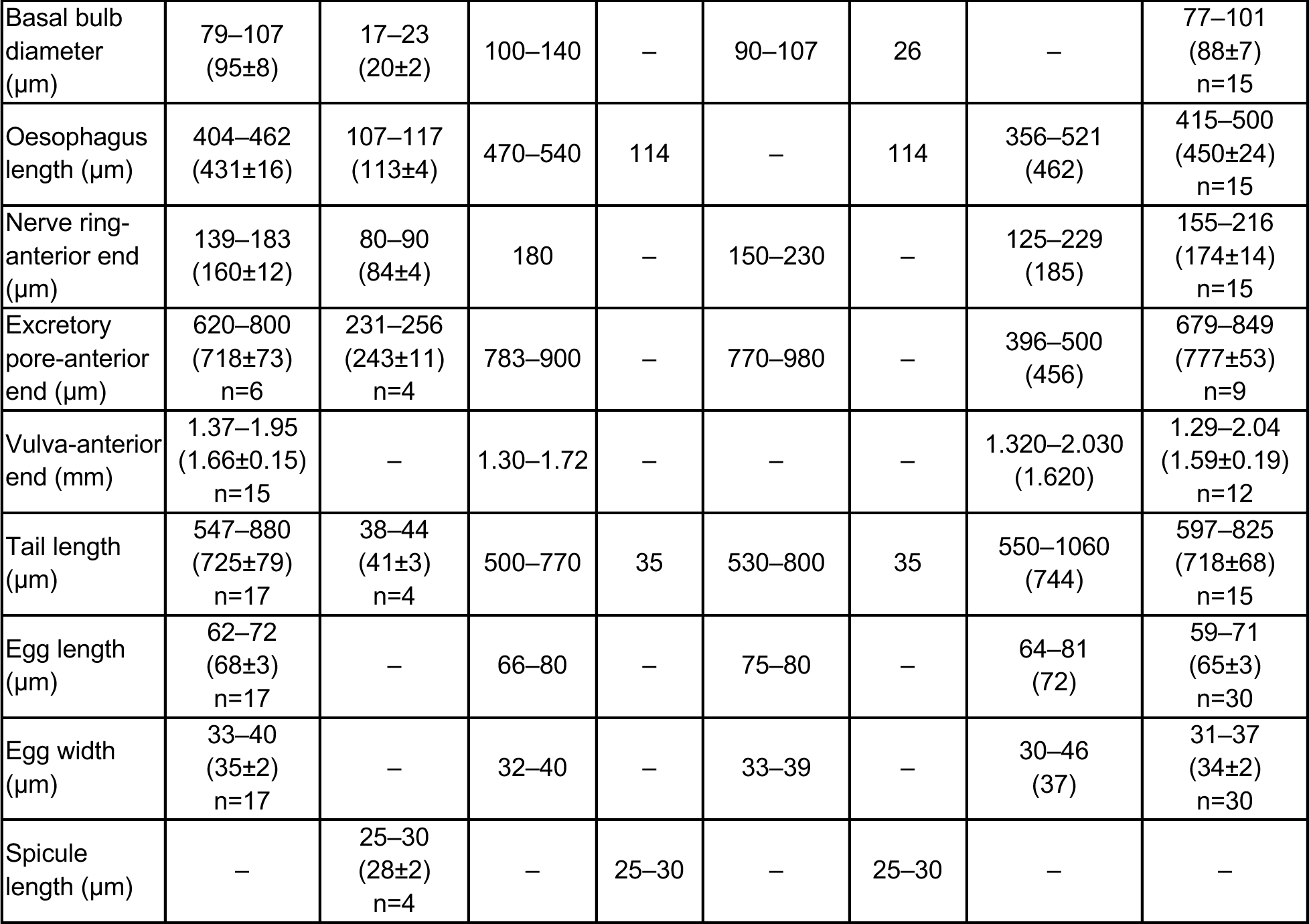
Comparison of the morphometrics of *Cranifera cranifera* (Chitwood, 1932) from captive-reared *Blaptica dubia* Serville, 1839 from Sweden with previous records of the same species. All the measurements are in the form of the range followed by mean plus standard deviation in parentheses, the number of measurements is also given if different from the total number of studied individuals.

| Locality | Sweden |  | Florida, USA |  | Germany |  | Costa Rica | Cuba |
| --- | --- | --- | --- | --- | --- | --- | --- | --- |
| Host | Captive colony of *Blaptica dubia* |  | Wild populations of *Blaberus cranifer*, *B. atropos* |  | Captive colony of *Blaptica dubia* |  | Wild populations of *Archimandrita tessellata* | Wild populations of *Blaberus discoidalis* |
| Authors | Current study |  | Chitwood (1932) |  | Leibersperger (1960) |  | Carreno & Tuhela (2003) | Morffe *et al.* (2022) |
| Character | females | males | females | males | females | males | females | females |
| # | 18 | 5 |  |  |  |  | 25 | 15 |
| a | 11.5–14.8<br>(13.1±1.1)<br>n=16 | 11.9–14.3<br>(13.1±1.0)<br>n=4 | – | – | – | 19.8 | – | 15.0–20.5<br>(16.8±1.5)<br>n=15 |
| b | 7.4–9.5<br>(8.8±0.7)<br>n=17 | 7.7–8.6<br>(8.1±0.4)<br>n=4 | – | – | – | 6.9 | – | 7.5–9.1<br>(8.1±0.5)<br>n=15 |
| c | 4.4–6.0<br>(5.2±0.5)<br>n=17 | 19.8–25.2<br>(22.6±2.2)<br>n=4 | – | – | – | 22.6 | – | 4.5–5.9<br>(5.1±0.4)<br>n=15 |
| V% | 40.9–47.3<br>(43.3±2.1)<br>n=15 | – | – | – | – | – | – | 41.7–48.1<br>(43.5±2.19)<br>n=12 |
| Total length (mm) | 2.9–4.2<br>(3.8±0.3)<br>n=17 | 0.87–0.96<br>(0.91±0.03)<br>n=4 | 3.2–3.8 | 0.79 | 3.5–4.9 | 0.79 | 2.8–6.7<br>(3.8) | 3.1–4.3<br>(3.6±0.3)<br>n=15 |
| Maximum width (µm) | 200–338<br>(289±41)<br>n=17 | 64–76<br>(69±5) | 200–250 | 40 | 190–290 | 40 | 172–400<br>(253) | 172–263<br>(219±30)<br>n=15 |
| First cephalic annule height (µm) | 7–12<br>(10±2)<br>n=12 | – | – | – | – | – | 8–11<br>(9) | 5–10<br>(7±2)<br>n=15 |
| First cephalic annule width (µm) | 35–40<br>(37±2)<br>n=12 | – | – | – | – | – |  | 26–36<br>(32±3)<br>n=15 |
| Stoma length (µm) | 17–26<br>(20±3)<br>n=15 | – | – | – | – | – | 17–28×27–32<br>(22×30) | 11–20<br>(15±3)<br>n=15 |
| Corpus length (µm) | 283–321<br>(304±12) | 49–55<br>(53±3) | – | – | – | – | 280–383<br>(340) | 296–369<br>(325±19)<br>n=15 |
| Isthmus length (µm) | 28–35<br>(31±3) | 34–39<br>(37±2) | 20–48 | 33 | 29–32 | 33 | 19–46<br>(32) | 25–36<br>(30±3)<br>n=15 |
| Basal bulb diameter (μm) | 79–107<br>(95±8) | 17–23<br>(20±2) | 100–140 | – | 90–107 | 26 | – | 77–101<br>(88±7)<br>n=15 |
| Oesophagus length (μm) | 404–462<br>(431±16) | 107–117<br>(113±4) | 470–540 | 114 | – | 114 | 356–521<br>(462) | 415–500<br>(450±24)<br>n=15 |
| Nerve ring-anterior end (μm) | 139–183<br>(160±12) | 80–90<br>(84±4) | 180 | – | 150–230 | – | 125–229<br>(185) | 155–216<br>(174±14)<br>n=15 |
| Excretory pore-anterior end (μm) | 620–800<br>(718±73)<br>n=6 | 231–256<br>(243±11)<br>n=4 | 783–900 | – | 770–980 | – | 396–500<br>(456) | 679–849<br>(777±53)<br>n=9 |
| Vulva-anterior end (mm) | 1.37–1.95<br>(1.66±0.15)<br>n=15 | – | 1.30–1.72 | – | – | – | 1.320–2.030<br>(1.620) | 1.29–2.04<br>(1.59±0.19)<br>n=12 |
| Tail length (μm) | 547–880<br>(725±79)<br>n=17 | 38–44<br>(41±3)<br>n=4 | 500–770 | 35 | 530–800 | 35 | 550–1060<br>(744) | 597–825<br>(718±68)<br>n=15 |
| Egg length (μm) | 62–72<br>(68±3)<br>n=17 | – | 66–80 | – | 75–80 | – | 64–81<br>(72) | 59–71<br>(65±3)<br>n=30 |
| Egg width (μm) | 33–40<br>(35±2)<br>n=17 | – | 32–40 | – | 33–39 | – | 30–46<br>(37) | 31–37<br>(34±2)<br>n=30 |
| Spicule length (μm) | – | 25–30<br>(28±2)<br>n=4 | – | 25–30 | – | 25–30 | – | – |

Males of the present population are longer (0.873–0.956 mm *vs.* 0.790 mm *vs.* 0.790 mm) and wider (64–76 μm *vs.* 40 μm *vs.* 40 μm) than those reported by Chitwood (1932) and Leibersperger (1960) respectively (Table 1). Their tail is also longer (38–44 μm *vs.* 35 μm *vs.* 35 μm). Compared with the record of Leibersperger (1960), the body of present male specimens is more robust (a = 11.9–14.3 *vs.* 19.8) and the relative length of the oesophagus is shorter (b = 7.7–8.5 *vs.* 6.9). Again, these differences can be equally attributed to both intraspecific variability of distant populations and measuring errors.

### Notes on female morphology

Females from the currently studied population are morphologically identical to the specimens from the original description (Chitwood, 1932). Most of the morphometrics also overlap (Table 1), and only the oesophagus length is slightly shorter on the present specimens (404–462 μm *vs.* 470–540 μm). The morphology and measurements of the individuals from the population described by Leibersperger (1960) also from *B. dubia* are also similar, except for the length of the eggs which are slightly shorter in current specimens (62–72 μm *vs.* 75–80 μm).

The excretory pore is more anterior in the specimens recorded by Carreno and Tuhela (2003) from Costa Rican *A. tessellata* (396–500 μm *vs.* 620–800 μm). The body of the specimens found in *B. discoidalis* from Cuba (Morffe et al., 2022) is slightly less robust (a = 15.0–20.5 *vs.* 11.5–14.8). All these differences can be equally attributed to both intraspecific variability of distant populations and measuring errors.

### Nuclear genome assembly

The final decontaminated genome assembly has a total length of 242 Mb, consisting of 7562 contigs, with an N50 = 43 kb and the longest contig being 1.10 Mb (Table 2). *BUSCO v.6.0.0* (Tegenfeldt et al., 2025) analysis using four latest reference datasets: *nematoda_odb12* (596 single copy-orthologs)*, nematoda_odb12.1* (1477 single-copy orthologs)*, chromadorea_odb12* (1126 single-copy orthologs) and *chromadorea_odb12.1* (2034 single-copy orthologs) identified 90.5-94% of the expected genes (Table 2), depending on the reference database used. The snail plot in Figure 3A. summarises the scaffold length distribution for the final assembly, while the blob plot in Figure 3B shows the distribution of contigs by GC proportion and coverage. High fragmentation of the final assembly and high rate of duplications after purging can be explained by population level heterozygosity of the input sample. Annotation of protein coding genes was not attempted due to absence of RNAseq data.

**Figure 3.**
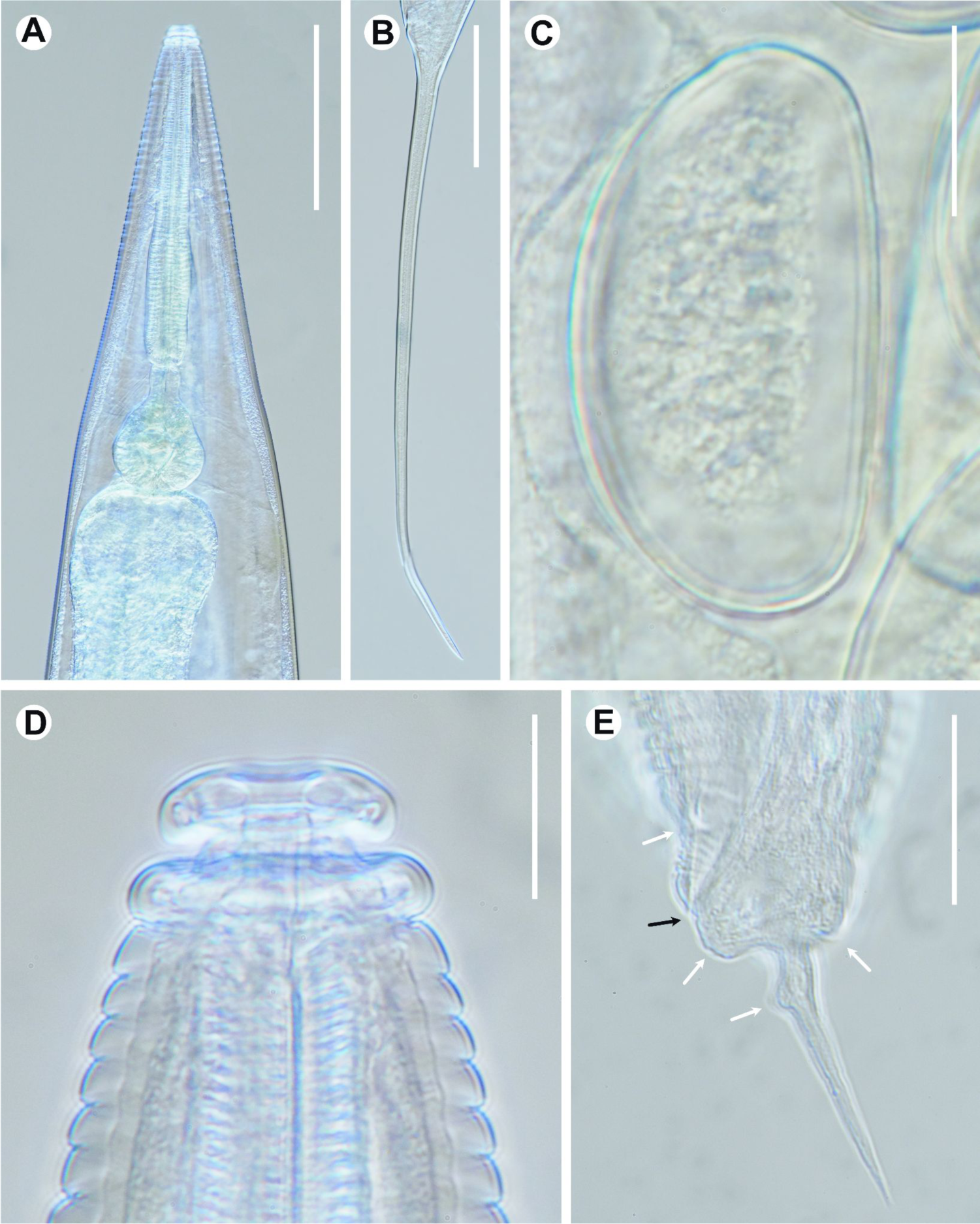
Light micrographs of a female (A-D) and a male (E) of *Cranifera cranifera* (Chitwood, 1932). A: Anterior body end showing pharyngeal region (ventral side to the right); B: Tail A: (ventral side to the left); C: Intra-uterine egg; D: Labial region; E: Caudal region (ventral side to the left), white arrows point to genital sensilla pairs, black arrow points to cloaca. Scale bars: A, B = 200 *μ*m. C, D, E = 25 *μ*m.

**Figure 4.**
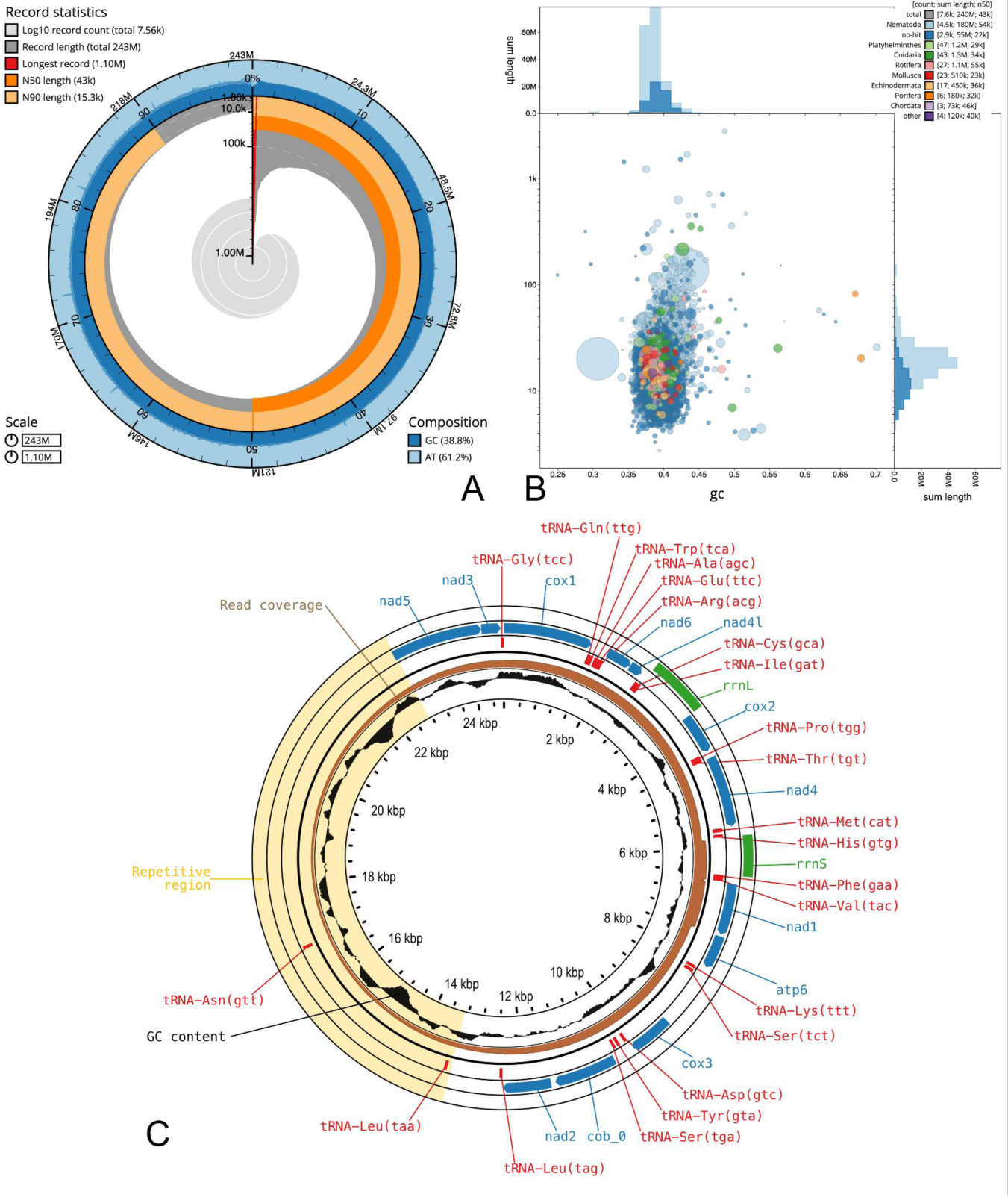
Nuclear (A-B) and mitochondrial (C) genomes of *Cranifera cranifera* (Chitwood, 1932). A: The BlobToolKit snail plot visualising the assembly metrics, the circumference represents the length of the whole genome sequence, the outermost blue tracks display the distribution of GC, AT, and N percentages, scaffolds are arranged clockwise from longest to shortest and are depicted in dark grey, the longest scaffold is indicated by the red arc, and the deeper orange and pale orange arcs represent the N50 and N90 lengths; B: The BlobToolKit blob plot visualising sequence coverage (vertical axis) and GC content (horizontal axis) with the circles representing scaffolds, with the size proportional to scaffold length and the colour representing phylum while the histograms along the axes display the total length of sequences distributed across different levels of coverage and GC content; C: Annotated mitogenome showing protein coding genes (blue), ribosomal RNA genes (green), transfer RNA genes (red), read coverage (brown), GC content (black) and repetitive region (yellow).

**Table 2.** Nuclear genome assembly statistics.

|  |  |  |  |  |
| --- | --- | --- | --- | --- |
| Accession | PRJEB115782 |  |  |  |
| Biosample | SAMEA122978875 |  |  |  |
| Assembly level | contig |  |  |  |
| Span | 242.73 Mb |  |  |  |
| Number of contigs | 7562 |  |  |  |
| Contig N50 | 43 kb |  |  |  |
| BUSCO reference database:<br>(# of single copy orthologs) | nematoda<br>odb12<br>(596) | nematoda<br>odb12.1<br>(1477) | chromadorea<br>odb12<br>(1126) | chromadorea<br>odb12.1<br>(2034) |
| Complete (all) | 542 (90.9%) | 1314 (89.0%) | 987 (87.7%) | 1763 (86.7%) |
| Complete (single-copy) | 460 (77.2%) | 1089 (73.7%) | 827 (73.4%) | 1491 (73.3%) |
| Complete (duplicated) | 82 (13.8%) | 225 (15.2%) | 160 (14.2%) | 272 (13.4%) |
| Fragmented | 18 (3.0%) | 47 (3.2%) | 42 (3.7%) | 78 (3.8%) |
| Missing | 36 (6.0%) | 116 (7.9%) | 97 (8.6%) | 193 (9.5%) |

### Mitochondrial genome assembly

The complete mitochondrial genome has a total length of 24644 bases (Figure 3C), including a region with high repeat content between 12653 and 22383 bases: repeat #1 is the largest duplicated block 2975-2977 bases long (12653–15626 and 19407–22383 positions); repeats #2 and #3 sit inside that same overall region; first copy of a 1318 bases long repeat #2 (15109–16426 positions) overlaps the tail of repeat #1’s first copy while the second copy of repeat #2 (17261–18578 positions) falls in the gap between repeat #1’s two copies; first copy of a 688 bases repeat #3 (17091–17778 positions) falls in the gap between repeat #1’s two copies while the second copy of repeat #3 (19237–19924 positions) partially overlaps with the second copy of repeat #1 (Figure 3C).

Annotation using both *Mitos2* (Bernt et al., 2013) and *Aragorn* (Laslett et al., 2004) identified a complete set of 12 protein-coding genes (atp6, cob, cox1-3, nad1-6, and nad4L) with atp8 being absent, as expected for nematodes, two rRNA genes and 22 tRNA genes, some of which were annotated several times by both *Mitos2* and *Aragorn*. Specifically, original annotation included eight tRNA-Lys(ttt) and four tRNA-Asn(gtt) copies, duplicates of tRNA-Ser(tct) and tRNA-Leu(taa), and a spurious tRNA-Tyr(gta) nested inside tRNA-Gly(tcc), both rRNAs were truncated (rrnS at 386 nt and rrnL at 666 nt), while several protein-coding genes had various annotation errors. The curated mitogenome includes the standard set of 22 tRNAs, with one functional copy kept for each tRNA duplicated in the repeat array, and rRNAs are re-annotated to 677 nt in rrnS and 961 nt in rrnL, in agreement with conserved motifs and gene lengths in the reference genomes. All protein coding genes have all start and stop codons in frame with no internal stops and their gene boundaries agree within a few amino acids with 13 published reference mitogenomes.

### Intragenomic variation of rRNA

Since 28S rRNA gene is the most commonly used barcode for species delimitation in Thelastomatidae, the newly assembled nuclear genome was mined for rRNA amplicon sequences in order to access intragenomic and intraspecific variation. Two amplicon variants of the 28S rDNA were mined from the newly assembled genome (Fig. 5A). In the Maximum Likelihood phylogram based on the 753 bp long alignment of the D2-D3 expansion domains of the 28S rDNA, one of these sequences (QB046965) forms a clade with the other three sequences of *C. cranifera* available in NCBI GenBank, although bootstrap support for this clade is moderate, and the internal topology is not resolved. The second copy of the gene (QB046966) was recovered as a sister clade to all other sequences. Based on the same alignment, the sequence of *C. cranifera* from *B. discoidalis* from Cuba (ON616698) was identical to QB046965. The sequence QB046966 differs in eight homologous positions from QB046965 (0.67% of genetic distance) and in five and eight homologous positions with the sequences from *C. cranifera* from the Cuban *Blaberus discoidalis* and from captive *Blaptica dubia* from Russia respectively (0.67% and 1.08% genetic distance, respectively). Seven of the eight variable bases between QB046965 and QB046966 are located within a variable segment 30 bases long and two of these correspond to indels (Fig. 5A).

**Figure 5.**
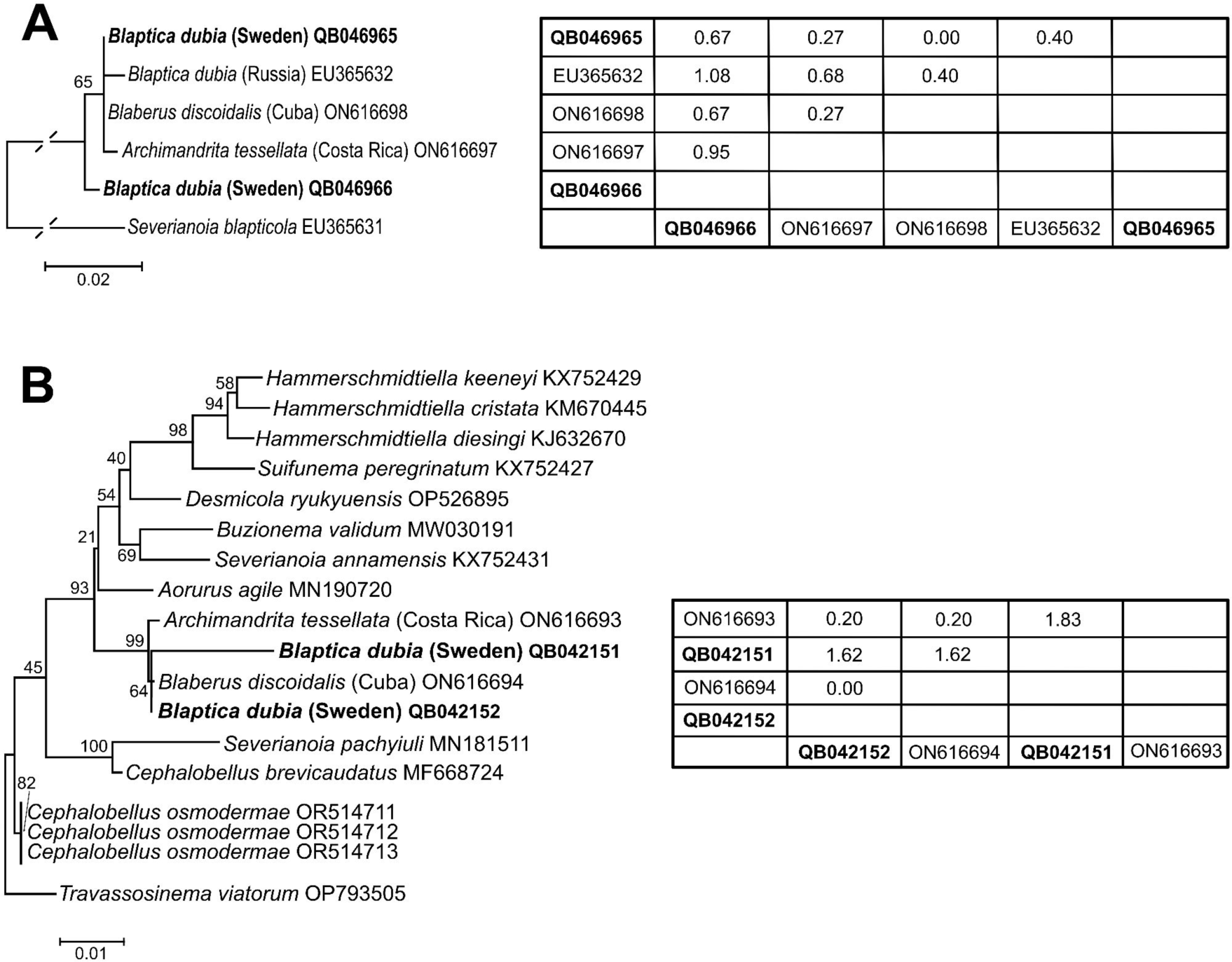
Maximum likelihood (ML) trees inferred from the D2-D3 28S rDNA for several sequences of *Cranifera cranifera* (Chitwood, 1932) with *Severianoia blapticola* Guzeeva, 2009 used as an outgroup (A) and the 18S rDNA for several sequences of *C. cranifera* and species of the family Thelastomatidae with *Travassosinema viatorum* Morffe, García & Hasegawa, 2023 used as an outgroup (B). Pairwise comparison of the *C. cranifera* sequences in the right. Sequences from the present study are highlighted in bold.

Despite the 18S rDNA gene being more useful at supraspecific level than as a species delimitation marker, it was included in the analyses. Two amplicon variants of the 18S rDNA were also recovered from the newly assembled genome (Fig. 5B) and used for the phylogenetic analysis together with the available sequences of *C. cranifera* and selected species of the family Thelastomatidae. In the Maximum Likelihood phylogram based on the 1508 bp long alignment all the sequences of *C. cranifera* form a strongly supported clade, with the sequence from *A. tessellata* from Costa Rica (ON616693) located in a basal position. The internal topology of this clade is not resolved, with the current sequences (QB042151 and QB042152) forming a clade with moderate support with the sequence of *C. cranifera* from *B. discoidalis* from Cuba (ON616698). Based on the same alignment, one of the current sequences (QB042152) was identical to the one from the Cuban *B. discoidalis*, similar to 28S (see above). The sequence QB042151 differs in 24 homologous positions (1.62% of genetic distance) with QB042152 and in 27 and 24 homologous positions with the sequences from *A. tessellata* from Costa Rica and *B. discoidalis* from Cuba (1.83% and 1.62% genetic distance, respectively) (Fig. 5B).

## Discussion

*Cranifera cranifera* is a common intestinal endobiont of several blaberid cockroaches, three of which are reared in captivity as pets (*Archimandrita tessellata* and *Blaberus craniifer*) or food source for other animals (*Blaptica dubia*), which makes this species easily accessible for both morphological and molecular studies. Despite this, many aspects of its biology remain poorly understood or completely unknown. For example, in addition to the present work, males of *C. cranifera* have only been previously recorded twice: first time in the original description of the species by Chitwood (1932) and second time in subsequent redescription by Leibersperger (1960). These authors only included drawings of the posterior end of the males and not from the anterior region and oesophagus, notwithstanding remarkable sexual dimorphism in this species (and in all Thelastomatoidea). Here, the anterior and oesophageal regions of the male are illustrated for the first time. The morphology of the oesophagus was described differently by Chitwood (1932) and Leibersperger (1960). Both considered the corpus as sub-divided in an anterior region [“a weak pseudobulb” according to Leibersperger (1960)] and a posterior region but did not describe their shape. Also, they considered the isthmus as short. In the present work we observed that this anterior region is the proper corpus and the posterior one constitutes a long isthmus. The arrangement of the copulatory papilliform sensilla observed by Chitwood (1932) and Leibersperger (1960) coincides with the current specimens: a pre-cloacal pair near the cloaca, and three post-cloacal pairs, one ad-anal pair, one sub-dorsal near the base of the tail appendage and one ventral, minute pair along the tail appendage. However, in the original description by Chitwood (1932) mentioned the copulatory papillae pattern as “caudal papillae consisting of one pair of preanal papillae, one pair of ad-anal or postanal papillae, and a median papilla near the base of the spinelike tail”, while the line drawing of the tail region showed both the ad-anal and the post-anal pairs of papillae. The latter points to a possible *lapsus calami* by the author.

The nuclear genomic resources for arthropod pinworms of the superfamily Thelastomatoidea are extremely limited (Lü et al., 2026), despite these species often being considered important ecological (not phylogenetical) intermediates between free-living nematodes and obligate endoparasites inhabiting some of the earliest terrestrial macrofauna (Sudhaus, 2010). Phylogenetically, they are nested within the more specialised groups of parasites (Sudhaus, 2010; Lü et al., 2026) which makes the origin and evolution of parasitism in Clade 3 nematodes an interesting puzzle. The nuclear genome of *C. cranifera* is the third for Thelastomatoidea arthropod pinworms, in addition to *Thelastoma shijiazhuangense* Zeng, Yuan, Liu & Meng, 2024 and *Travassosinema thyropygi* Hunt, 1996 (Lü et al., 2026), the latter two being based on Illumina short read sequencing with no assembly or annotation files being publicly available. As such, building a comprehensive and dependable phylogenetic framework for early branching Clade 3 nematodes remains nearly impossible, due to huge gaps in taxonomic sampling for sequencing projects.

While there are many more mitochondrial genomes available for Clade 3 nematodes, including Thelastomatoidea (Lü et al., 2026; Xu et al. 2026), their utility for phylogenetic analysis remains questionable, mainly due to topological discrepancies with phylogenies based on nuclear genomes in the interrelationships between Clade 3 (Spirurina) and Clade 4 (Tylenchina) nematodes (Zhan et al., 2026). Gene arrangement in mitochondrial genomes also vary considerably across the entire Nematoda, being relatively conserved in some lineages and showing no conserved pattern at all in other groups (Du et al., 2022; Nikolaeva et al., 2025; Lü et al., 2026; Tomé et al., 2026), which can be partially explained by sequencing bias: conserved patterns are often observed in closely related species, members of the same genus, while more diverse arrangement of protein coding genes is more commonly observed in sparsely sequenced lineages. As such, we are not discussing the variations of gene order in mitochondrial genomes of nematodes, awaiting for a much larger body of information. Instead, we would like to highlight the impact of repetitive regions on the assembly and annotation of nematode mitochondrial genomes. The mitochondrial genome of *C. cranifera* is 24644 bases long including a region with complex repeat content spanning 9729 bases, with the largest repeat fragment being almost 3000 bases long. This region was assembled both by *MitoHifi* and our custom pipeline. We can not completely exclude the possibility that it is an assembly artifact, since alternative assemblies 17888, 22497 and 31115 bases long were also recovered.

However, mapping raw reads against all four alternative assemblies revealed assembly collapse in both 17888 and 22497 bases long assemblies, and coverage drop in 31115 bases long assembly. Similarly sized non-coding regions were identified in mitogenomes of other nematode species as well, specifically in *Hoplolaimus columbus* (7661 bp and 3157 bp long), *Pratylenchus vulnus* (6847 bp), *Meloidogyne chitwoodi* (5404 bp), *M. graminicola* (5063 bp) and *Meloidogyne incognita* (4097 bp) (Sultana et al., 2013; Humphreys-Pereira and Elling, 2014; Sun et al., 2014; Ma et al., 2020). Therefore, presence of duplicated regions in mitogenome assemblies based on short read data (where duplications are likely to be collapsed during assembly) may not be recovered, unless read coverage is used for verification.

Close match between recovered sequences of 28S rDNA and 18S rDNA (especially QB046965 and QB042152, respectively) and reference D2-D3 domains of the 28S rDNA and 18S rDNA available from the NCBI GenBank confirms the identity of the studied population as *C. cranifera* and complements the morphological and morphometric data. The basal position of the QB046966 in the 28S rDNA phylogram compared to all other sequences of *C. cranifera* in our opinion represents intragenomic variation, a less common sequence variant. Despite the fact that QB042151 and QB042152 are located in the same clade of the 18S rDNA tree, the divergence rate of QB042151 also points to the presence of intragenomic variation in this gene. Intragenomic variation in rRNA sequences has been previously documented in several groups of free-living and parasitic nematodes where several authors (*i.e.* Bik et al., 2013; Qing et al., 2019), where one of the sequence variants can be more divergent from the other, creating a polyphyletic topology (Fig. 5A) and supporting the hypothesis that such divergent variants are less common copies, negatively selected during regular PCR amplification, but readily identifiable in whole-genome datasets, and should be accepted as a fact of intragenomic heterogeneity of the rRNA operon and not a signal of interspecific divergence of cryptic speciation.

Last but not least, the actual impact of thelastomatid nematodes on their hosts remains poorly understood. Studies suggest that thelastomatid nematodes specialise on feeding on the gut content of detritivorous host (Hominick and Davey, 1973) and may positively affect the diversity of the host microbiome (Vicente et al., 2016), but little is known on how this affects the host health (McCallister, 1988), and nothing about whether the host provides a response. Being relatively easy to identify comparing to other Thelastomatoidea, having a simple life cycle, inhabiting commercially accessible and easy to maintain hosts that are currently completely exempt from ethical review processes, being more closely related to many parasites of veterinarian and human health importance than *C. elegans*, and having genomic resources available now, *C. cranifera* has a great potential to be used as a model system to study not only the evolution of parasitism in general but also the acquisition of anthelminthic resistance and how to counteract it. Other species from the same family have already been used to study anthelmintic response in the past (Holoman, 1980; Yu and Crites, 1990).

## Data availability

Morphological vouchers on permanent glass slides are deposited in the invertebrate collection of the Department of Zoology, Swedish Museum of Natural History (SMNH), Stockholm, Sweden (accession numbers SMNH 231508–231518). Raw data and assembled nuclear genome assembly are available from the European Nucleotide Archive under the accession number PRJEB115782. Raw reads under BioSample number SAMEA122978875 and accession number ERR17501394 respectively. The curated nuclear genome is available under the accession number ERZ30225045 and at https://www.ebi.ac.uk/ena/browser/view/GCA_986284995. The mitochondrial genome is available from the NCBI GenBank under the accession number QB100924.

## Author contribution

Jans Morffe performed collection and preservation of nematodes and DNA extraction and morphological analysis; Nadège Guiglielmoni, Nadim Wassey, Karim Gueddach, Anja Schuster, and Kerstin Becker generated genomic data; Jens Bast and Philipp Schiffer acquired funding and oversaw generation of genomic data; Oleksandr Holovachov conceptualised and supervised the study in general as well as performed genome assemblies. All authors contributed to writing and approved the final version of the manuscript.

## Acknowledgements

The authors are grateful to Niklas Apelqvist (Swedish Museum of Natural History) for providing the *Blaptica dubia* colony. Computational support and infrastructure were provided by the “Centre for Information and Media Technology” (ZIM) at the University of Düsseldorf (Germany).

## Ethics approval and consent to participate

Not applicable. No ethics approval is necessary.

## References

Adamson ML and Van Waerebeke D (1992) Revision of the Thelastomatoidea, Oxyurida of invertebrate hosts I. Thelastomatidae. Systematic Parasitology 21, 21–63. 10.1007/BF00009911

Allio R, Schomaker-Bastos A, Romiguier J, Prosdocimi F, Nabholz B and Delsuc F (2020) MitoFinder: Efficient automated large-scale extraction of mitogenomic data in target enrichment phylogenomics. Molecular Ecology Resources 20, 892–905. 10.1111/1755-0998.13160

Altschul SF, Gish W, Miller W, Myers EW and Lipman DJ. (1990) Basic local alignment search tool. Journal of Molecular Biology 215, 403–410. 10.1016/S0022-2836(05)80360-2

Bernt M, Donath A, Jühling F, Externbrink F, Florentz C, Fritzsch G, Pütz J, Middendorf M and Stadler PF (2013) MITOS: Improved de novo metazoan mitochondrial genome annotation. Molecular Phylogenetics and Evolution 69, 313–319. 10.1016/j.ympev.2012.08.023

Bik, HM, Fournier D, Sung W, Bergeron RD and Thomas WK (2013) Intra-Genomic Variation in the Ribosomal Repeats of Nematodes. PLoS ONE 8, e78230. 10.1371/journal.pone.0078230

Buchfink B, Reuter K and Drost HG (2021) Sensitive protein alignments at tree-of-life scale using DIAMOND. Nature Methods 18, 366–368. 10.1038/s41592-021-01101-x

Camino NB and Achinelly MF (2012) A new species of the genus *Cranifera* Kloss, 1960 (Thelastomatidae, Nematoda) parasitizing larvae of Scarabaeidae (Coleoptera) from Argentina. Estudos de Biologia 34, 57–59. 10.7213/estud.biol.6124

Carreno RA (2014) The systematics and evolution of pinworms (Nematoda: Oxyurida: Thelastomatoidea) from invertebrates. Journal of Parasitology 100, 553–560. 10.1645/14-529.1

Carreno RA and Tuhela L (2011) Thelastomatid nematodes (Oxyurida: Thelastomatoidea) from the peppered cockroach, *Archimandrita tessellata* (Insecta: Blattaria) in Costa Rica. Comparative Parasitology 78, 39–55.

Chakraborty M, Baldwin-Brown JG, Long AD and Emerson JJ (2016) Contiguous and accurate de novo assembly of metazoan genomes with modest long read coverage. Nucleic Acids Research 44, e147. 10.1093/nar/gkw654

Challis R, Richards E, Rajan J, Cochrane G and Blaxter M (2020) BlobToolKit – interactive quality assessment of genome assemblies. G3 Genes|Genomes|Genetics 10, 1361–1374. 10.1534/g3.119.400908

Chen S (2023) Ultrafast one-pass FASTQ data preprocessing, quality control, and deduplication using fastp. iMeta 2, e107. 10.1002/imt2.107

Cheng H, Asri M, Lucas J, Koren S and Li H (2024) Scalable telomere-to-telomere assembly for diploid and polyploid genomes with double graph. Nature Methods 21, 967–970. 10.1038/s41592-024-02269-8

Chitwood BG (1932) A synopsys of the nematodes parasitic in insects of the family Blattidae. Zeitschrift für Parasitenkunde 5, 14–50.

Coy A and García N (1995) Nuevas especies de nemátodos parásitos de insectos mexicanos. AvaCient 12, 10–15.

Danecek P, Bonfield JK, Liddle J, Marshall J, Ohan V, Pollard MO, Whitwham A, Keane T, McCarthy SA, Davies RM and Li H (2021) Twelve years of SAMtools and BCFtools. GigaScience 10, giab008. 10.1093/gigascience/giab008

De Coster W and Rademakers R (2023) NanoPack2: population-scale evaluation of long-read sequencing data, Bioinformatics 39, btad311. 10.1093/bioinformatics/btad311

Devi MP, Haryoso E, Rais EI, Karuniawan A, Yahya MQ, Richaud A, Wang J, Rockman MV, Tarno H and Félix M-A (2025) Five new *Caenorhabditis* species from Indonesia provide exceptions to Haldane’s rule and partial fertility of interspecific hybrids. G3 Genes|Genomes|Genetics 15, jkaf134. 10.1093/g3journal/jkaf134

Du H, Guo F, Gao Y, Wang X, Qing X and Li H (2022) Complete mitogenome of *Cruznema tripartitum* confirms highly conserved gene arrangement within Family Rhabditidae. Journal of Nematology 54, 1–10. 10.2478/jofnem-2022-0029

Dulovic A, Renahan T, Röseler W, Rödelsperger C, Rose AM and Streit A (2020) *Rhabditophanes diutinus* a parthenogenetic clade IV nematode with dauer larvae. PLoS Pathogens 16: e1009113. 10.1371/journal.ppat.1009113

Edgar RC (2004) MUSCLE: multiple sequence alignment with high accuracy and high throughput. Nucleic Acids Research 32, 1792–1797. 10.1093/nar/gkh340

Gendron EM, Sevigny JL, Byiringiro I, Thomas WK, Powers TO and Porazinska DL (2023) Nematode mitochondrial metagenomics: a new tool for biodiversity analysis. Molecular Ecology Resources 23, 975–989. 10.1111/1755-0998.13761

Grant JR, Enns E, Marinier E, Mandal A, Herman EK, Chen C, Graham M, Van Domselaar G and Stothard P (2023) Proksee: in-depth characterization and visualization of bacterial genomes. Nucleic Acids Research 51(W1), W484–W492. 10.1093/nar/gkad326

Guan D, McCarthy SA, Wood J, Howe K, Wang Y and Durbin R (2020) Identifying and removing haplotypic duplication in primary genome assemblies. Bioinformatics 36, 2896–2898. 10.1093/bioinformatics/btaa025

Holoman VL (1980) A study of oxyuroid nematode feeding behavior and the use of cockroaches as an insect model for testing anthelmintics. The Ohio State University, Columbus.

Hominick WM and Davey KG (1973) Food and the spatial distribution of adult female pinworms parasitic in the hindgut of *Periplaneta americana* L. International Journal of Parasitology 3, 759–771. 10.1016/0020-7519(73)90067-2

Humphreys-Pereira DA and Elling AA (2014) Mitochondrial genomes of *Meloidogyne chitwoodi* and *M. incognita* (Nematoda: Tylenchina): Comparative analysis, gene order and phylogenetic relationships with other nematodes. Molecular and Biochemical Parasitology 194, 20–32. 10.1016/j.molbiopara.2014.04.003

Kimura M (1980) A simple method for estimating evolutionary rate of base substitutions through comparative studies of nucleotide sequences. Journal of Molecular Evolution 16, 111–120. 10.1007/BF01731581

Kloss G (1960). Organizaçao filogenetica dos nematoides parasitos intestinais dos artropodes (Nota previa). Atas da Sociedade de Biologia do Rio de Janeiro 4, 51–55.

Kolmogorov M, Yuan J, Lin Y and Pevzner PA (2019) Assembly of long, error-prone reads using repeat graphs. Nature Biotechnology 37, 540–546. 10.1038/s41587-019-0072-8

Koren S, Walenz BP, Berlin K, Miller JR, Bergman NH and Phillippy AM (2017) Canu: scalable and accurate long-read assembly via adaptive k-mer weighting and repeat separation. Genome Research 27, 722–736. 10.1101/gr.215087.116

Kumar S, Stecher G, Suleski M, Sanderford M, Sharma S and Tamura K (2024) Molecular Evolutionary Genetics Analysis Version 12 for adaptive and green computing. Molecular Biology and Evolution 41, 1–9. 10.1093/molbev/msae263

Larsson A (2014) AliView: a fast and lightweight alignment viewer and editor for large datasets. Bioinformatics 30, 3276–3278. 10.1093/bioinformatics/btu531

Laslett D and Canback B (2004) ARAGORN, a program to detect tRNA genes and tmRNA genes in nucleotide sequences. Nucleic Acids Research 32, 11–16. 10.1093/nar/gkh152

Launay C, Félix MA, Dieng J and Delattre M (2020) Diversification and hybrid incompatibility in auto-pseudogamous species of *Mesorhabditis* nematodes. BMC Evolutionary Biology 20, 105. 10.1186/s12862-020-01665-w

Leibersperger E. (1960) Die Oxyuroidea der europaischen Arthropoden. Parasitologische Schriftenreihe 11, 150 pp.

Li H (2021) New strategies to improve minimap2 alignment accuracy. Bioinformatics 37, 4572– 4574. 10.1093/bioinformatics/btab705

Lü L, Zhang SJ, Gu XH, Zhang D, Hugot JP, Gibson DI, Chen HX, Zhang Y, Xu HR and Li L (2026) Phylogenomics and the evolutionary history of the Oxyurida (pinworms). Cladistics. 10.1111/cla.70047

Ma X, Agudelo P, Richards VP and Baeza JA (2020) The complete mitochondrial genome of the Columbia lance nematode, *Hoplolaimus columbus*, a major agricultural pathogen in North America. Parasites & Vectors 13, 321. 10.1186/s13071-020-04187-y

Malysheva SV, Shmatko VY and Spiridonov SE (2020) Revision of *Severianoia* (Schwenk, 1926) Travassos, 1929 (Nematoda: Oxyuridomorpha) with proposal of *S. pachyiuli* n. sp. from millipedes of the Western Caucasus. Nematology 23, 467–481. 10.1163/15685411-bja10053

McCallister GL (1988) The effect of *Thelastoma bulhoesi* and *Hammerschmidtiella diesingi* (Nematoda, Oxyurata) on host size and physiology in *Periplaneta americana* (Arthropoda, Blattidae). Proceedings of the Helminthological Society of Washington 55, 12–14.

Morffe J, García N, Véliz L, Hasegawa K and Carreno RA (2022) Morphological and molecular characterization of two species of nematodes (Oxyuridomorpha: Thelastomatoidea: Protrelloididae, Thelastomatidae) parasitic in the cockroach *Blaberus discoidalis* Serville (Blattaria: Blaberidae) from Cuba. Zootaxa 5194, 92–108. 10.11646/zootaxa.5194.1.5

Nie F, Ni P, Huang N, Zhang J, Wang Z, Xiao C, Luo F and Wang J (2024) De novo diploid genome assembly using long noisy reads. Nature Communications 15, 2964. 10.1038/s41467-024-47349-7

Nikolaeva OV, Ovcharenko AS, Khorkhordina PV, Miroliubova TS, Sadovskaya NS, Scobeyeva VA, Sanamyan NP, Panina EG, Mikhailov KV, Rusin LY, Tchesunov AV and Aleoshin VV (2025) Gene order in mitochondrial DNA affects the abundance of their transcripts (a case of marine nematodes). Biochemistry 90, 1843–1861. 10.7868/S3034529425110195

Pettrich LC, Suwanngam A, Guiglielmoni N, Ledoux N, Villegas LI, Treonis A, Stevens L, Kieninger M, Paulini M, Blaxter M, Waldvogel A-M, Holovachov O and Schiffer PH (2026) *Panagrolaimus einhardii* sp. nov. and two sisters of fortune. Zoological Journal of the Linnean Society 208: zlag125. 10.1093/zoolinnean/zlag125

Pryszcz LP and Gabaldón T (2016) Redundans: an assembly pipeline for highly heterozygous genomes. Nucleic Acids Research 44, e113. 10.1093/nar/gkw294

Qing X, Bik H, Yergalyev TM, Gu J, Fonderie P, Brown-Miyara S, Szitenberg A and Bert W (2020) Widespread prevalence but contrasting patterns of intragenomic rRNA polymorphisms in nematodes: Implications for phylogeny, species delimitation and life history inference. Molecular Ecology Resources 20, 318–332. 10.1111/1755-0998.13118

Röseler W, Yoshida K and Rödelsperger C (2025) Genome announcement: further improved genome assembly of *Parapristionchus giblindavisi*. Journal of Nematology 57, 1. 10.2478/jofnem-2025-0026

Seinhorst JW (1959) A rapid method for the transfer of nematodes from fixative to anhydrous glycerin. Nematologica 4, 67–69. 10.1163/187529259X00381

Shen W, Sipos B and Zhao L (2024) SeqKit2: a Swiss army knife for sequence and alignment processing. iMeta 3, e191. 10.1002/imt2.191

Slos D, Sudhaus W, Stevens L, Bert W and Blaxter M (2017) *Caenorhabditis monodelphis* sp. n.: defining the stem morphology and genomics of the genus *Caenorhabditis*. BMC Zoology 2, 4. 10.1186/s40850-017-0013-2

Spiridonov SE and Guzeeva EA. (2009) Phylogeny of nematodes of the superfamily Thelastomatoidea (Oxyurida) inferred from LSU rDNA sequence. Russian Journal of Nematology 17, 127–134.

Stevens L, Félix MA, Beltran T, Braendle C, Caurcel C, Fausett S, Fitch D, Frézal L, Gosse C, Kaur T, Kiontke K, Newton MD, Noble LM, Richaud A, Rockman MV, Sudhaus W and Blaxter M (2019) Comparative genomics of 10 new *Caenorhabditis* species. Evolutionary Letters 3, 217–236. 10.1002/evl3.110

Sudhaus W (2010) Preadaptive plateau in Rhabditida (Nematoda) allowed the repeated evolution of zooparasites, with an outlook on evolution of life cycles within Spiroascarida. Palaeodiversity 3 (supplement), 117–130.

Sultana T, Kim J, Lee SH, Han H, Kim S, Min GS, Nadler SA and Park JK (2013) Comparative analysis of complete mitochondrial genome sequences confirms independent origins of plant-parasitic nematodes. BMC Evolutionary Biology 13, 12. 10.1186/1471-2148-13-12

Sun L, Zhuo K, Lin B, Wang H and Liao J (2014) The complete mitochondrial genome of *Meloidogyne graminicola* (Tylenchina): a unique gene arrangement and its phylogenetic implications. PLoS ONE 9, e98558. 10.1371/journal.pone.0098558

Tegenfeldt F, Kuznetsov D, Manni M, Berkeley M, Zdobnov EM and Kriventseva EV (2025) OrthoDB and BUSCO update: annotation of orthologs with wider sampling of genomes. Nucleic Acids Research 53, D516–D522. 10.1093/nar/gkae987

The Galaxy Community (2026) The Galaxy platform for accessible, reproducible, and collaborative data analyses: 2024 update. Nucleic Acids Research 54, W105–W116. 10.1093/nar/gkag469

Tomé B, Harris DJ, Jorge F, de Sousa A, Archer J, Muñoz-Merida A, Rato C, Perera A, Mulder KP (2026) Complete mitochondrial genomes of four pharyngodonid nematodes reveal extensive gene order rearrangement. Journal of Helminthology 100, e32. 10.1017/S0022149X2610128X

Uliano-Silva M, Ferreira JGRN, Krasheninnikova K, Darwin Tree of Life Consortium, Formenti G, Abueg L, Torrance J, Myers EW, Durbin R, Blaxter M and McCarthy SA (2023) MitoHiFi: a python pipeline for mitochondrial genome assembly from PacBio high fidelity reads. BMC Bioinformatics 24, 288. 10.1186/s12859-023-05385-y

Vicente CS, Ozawa S and Hasegawa K (2016) Composition of the cockroach gut microbiome in the presence of parasitic nematodes. Microbes and Environments 31, 314–320. 10.1264/jsme2.ME16088

Walker BJ, Abeel T, Shea T, Priest M, Abouelliel A, Sakthikumar S, Cuomo CA, Zeng Q, Wortman J, Young SK and Earl AM (2014) Pilon: an integrated tool for comprehensive microbial variant detection and genome assembly improvement. PLoS ONE 9, e112963. 10.1371/journal.pone.0112963

Xu HR, Zhang Y, Zhang K and Li (2026) Revision and molecular phylogeny of the genus *Travassosinema* (Nematoda: Oxyurida: Travassosinematidae) from millipedes in China, with broad reference to the Clade III nematodes. Zoological Journal of the Linnean Society 207, zlag105. 10.1093/zoolinnean/zlag105

Yu X and Crites JL (1990) The effectiveness of two anthelmintics, Vermox (mebendazole) and Povan (pyrvinium pamoate), on thelastomatid nematodes (Nematoda: Oxyuroidea) of the cockroach, *Gromphadorhina portentosa*. Ohio Journal of Science 90, 152–155.

Zhan F, Shen C, Mundock IM, Ren Y, Li H, Xue Q and Guo W (2026) Draft whole-genome and mitochondrial genome assemblies of *Steinernema tarimense* and *Heterorhabditis* sp. XJ-55. Journal of Helminthology 100, e63. 10.1017/S0022149X26101667

